# Information use during movement regulates how fragmentation and loss of habitat affect body size

**DOI:** 10.1101/265025

**Authors:** Jasmijn Hillaert, Martijn L. Vandegehuchte, Thomas Hovestadt, Dries Bonte

## Abstract

An individual's body size is central to its behavior and physiology, and tightly linked to its dispersal ability. The spatial arrangement of resources and a consumer's capacity to locate them are therefore expected to exert strong selection on consumer body size.

We investigated the evolutionary impact of both the fragmentation and loss of habitat on consumer body size and its feedback effects on resource distribution, under varying levels of information use during the settlement phase of dispersal. We developed a mechanistic, individual-based, spatially explicit model, including several allometric rules for key consumer traits. Our model reveals that as resources become more fragmented and scarce, informed settlement selects for larger body sizes while random settlement promotes small sizes. Information use may thus be an overlooked explanation for the observed variation in body size responses to habitat fragmentation. Moreover, we find that resources can accumulate and aggregate if information on resource abundance is incomplete. Informed movement results in stable resource-consumer dynamics and controlled resources across space. However, habitat fragmentation and loss destabilize local dynamics and disturb resource suppression by the consumer. Considering information use during movement is thus critical to understand the eco-evolutionary dynamics underlying the functioning and structuring of consumer communities.

## Background

Habitat fragmentation and loss pose severe threats to size diversity at the population and community level, affecting size distributions. Eventually, shifts in size distributions impact ecosystem dynamics (incl. fluxes of nutrients) and functioning [1,2]. As such, a better understanding of the impact of habitat fragmentation and loss on body size distribution through selection is crucial [3].

An organism’s body size is one of its most comprehensive characteristics. Because of the %-scaling rule with metabolic rate, body size is strongly correlated with an array of functional traits, such as ingestion rate, movement speed and developmental time [4,5]. As such, body size represents the outcome of several selective pressures acting on different life history traits, setting boundaries to the ecology, physiology and functioning of an individual [4]. Body size distributions within communities additionally affect intra- and interspecific interactions, important higher-level properties of food webs, and ecosystem functioning [6–9]. Overall, body size can be considered a universal trait constraining ecological and evolutionary dynamics [10,11].

Body size distributions are strongly determined by the availability of resources and their distribution across space [12–17]. Hollings’ textural discontinuity hypothesis even states that the modes of a body size distribution reflect the foraging scales with the highest resource amounts [15–19]. As habitat fragmentation and destruction progress, the spatial distribution of resources is altered, yet the consequences for (future) body size distributions are unclear. On the one hand, large-bodied individuals may be selected as they have high starvation resistance and are able to cover large distances [4,20,21]. On the other hand, small-sized individuals may have the benefit of short developmental times and low energy requirements [4]. Empirical studies illustrate positive [22–24], negative [25,26] or insignificant [20] effects of fragmentation on average body size within populations. At the community level, shifts in species abundances and therefore size distributions strongly depend on taxonomical group [27]. Despite this variation in empirical results and the absence of a consensus in theoretical work, several theoretical studies have acknowledged a strong dependency of size distributions on habitat configuration [12,28-31]. Habitat fragmentation and loss are considered two distinct processes [32]. Habitat loss results in a decreased percentage of suitable habitat, whereas habitat fragmentation implies a decrease in its spatial autocorrelation [32]. Most experimental studies focus on their joint effect using the term ‘fragmentation’ or ‘landscape simplification’, without assessing the effects of each of these processes independently (e.g. [20,2427,33] (but see [23] for an exception). This is surprising as the effect of spatial autocorrelation is highest, and therefore most relevant, in landscapes with low percentages of suitable habitat [32]. Furthermore, most fragmentation studies focused on mammals and birds and were therefore performed at large spatial scales [34,35]. However, small spatial scales are most important for arthropods that do not disperse via the air or by flying [27,34]. Still, only few empirical studies have investigated changes in arthropod size distributions at such scales (e.g. [20], exception: [27]).

Not only resource availability, but also the type of movement and dispersal regulate how populations and communities are spatially structured [36]. High movement frequencies result in spatially coupled populations, whereas low frequencies result in classic metapopulations or -communities [37]. Furthermore, movement behavior not only depends on an individual’s body size but also on the information perceived during movement, which enables individuals to continuously update decisions on how far to move and where to settle [38]. The available information differs between organisms, depending on the complexity of their senses. As proven by theoretical studies, some degree of informed settlement already strongly affects spatial dynamics and coexistence compared to random walk [39,40]. Therefore, the effect of informed movement should be incorporated in studies focusing on movement ecology [41].

We designed an individual-based, spatially explicit model to study the effect of habitat configuration on the body size distribution of a population or community of arthropods. As the level of information perceived during movement is crucial for movement and therefore body size evolution, we investigated a possible interaction with this trait. We applied a mechanistic approach by incorporating established allometric rules linking body size with movement speed, movement costs, basal metabolic rate, ingestion rate, developmental time, and clutch size into our model. Due to the universal nature of these allometric rules, our conclusions on the effects of fine-grained fragmentation may apply to a wide range of taxa [5].

## Methods

The applied model is a spatially explicit, discrete-time model with overlapping generations. One time step corresponds to one day within the lifetime of the consumer. We here took an arthropod-centered approach and parameterized allometric rules for a haploid, parthenogenetic arthropod species feeding on plants (the resource), with a semelparous lifecycle. See table S1.1 for an overview of all parameters applied within the model.

### The landscape

The landscape is a cellular grid of 100 by 100 cells and is generated using the Python package NLMpy [42]. Each cell within the landscape has a side length (SL) of 0.25 m and therefore a total surface of 0.0625 m^2^. Within the landscape, a distinction is made between suitable and unsuitable habitat. Only within suitable habitat, the resource is able to grow. When testing the effect of landscape configuration, the proportion of suitable habitat (P) and habitat autocorrelation (H) were varied between landscapes. Habitat availability increases with P, whereas habitat fragmentation decreases with *H.* The following values were assigned to P: 0.05, 0.20, 0.50 or 0.90. *H* equaled either 1 (in all four cases), 0.5 (when *P* equaled 0.05 or 0.20) or 0 (when *P* equaled 0.05). As such, highly fragmented landscapes with a high amount of suitable habitat were not included in the analysis as these rarely occur in nature [43].

### The resource

Resources are not individually modeled but by a logistic growth model for each habitat cell. Local resource biomass is represented as the total energetic content of resource tissue within that cell *(R_x_,y* in joules). This resource availability grows logistically in time depending on the resource’s carrying capacity (K) and intrinsic growth rate (r). In any cell, a fixed amount of resource tissue *(E_nc_,* in Joules, fixed at *1 J)* is non-consumable by the consumer species, representing below-ground plant parts. As such, *Enc* is the minimum amount of resource tissue present within a suitable cell, even following local depletion by the consumer species.

### The consumer

All consumers are individually modelled within the landscape. The consumer has two life stages: a juvenile and adult life stage. Within a day, both stages have the chance to execute different events (see Figure 1).

**Figure 1:**
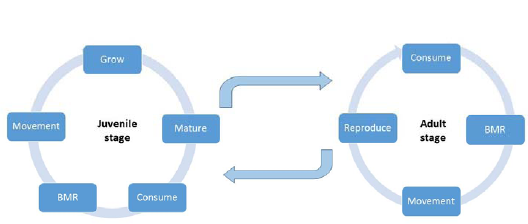
A comparison of daily events for the juvenile and adult stage of the consumer. BMR stands for the basal metabolic rate costs.

First, an individual nourishes its energy reserve by consumption. Second, the energy reserve is depleted by the cost of daily maintenance (i.e. basal metabolic rate) and the cost of movement. To assess the effect of informed settlement on our results, three different types of movement (see below) were implemented within the model. Third, juveniles may further deplete the energy reserve by growth, eventually resulting in maturation if they approximate their adult size *(W_max_).* Resources that were not utilized are stored within the energy reserve. Adults can only reproduce if their internally stored energy *(E_r_)* exceeds a predefined amount. As the consumer species is semelparous, adults die after reproduction. How body size affects each of these events is explained in supplementary material part 1.

Individual body size at maturity *(W_max_,* in kg) is coded by a single gene. Adult size is heritable and may mutate with a probability of 0.001 during reproduction. A new mutation is drawn from the uniform distribution *[W_max_* - (W_max_2), *W_max_* + *(W_max_/2)]* with *W_max_* referring to the adult size of the parent. New mutations may not exceed the predefined boundaries [0.01g, 3g] that represent absolute physiological limits. As such, our minimum adult size corresponds to the size of a small grasshopper such as *Tetrix undulata* (0.01 g) and the maximum size (3 g) to that of some longhorn beetles (Cerambycidae), darkling beetles (Tenebrionidae), scarab beetles (Scarabaeidae) or grasshoppers (Acrididae). New variants of this trait may also originate by immigration (see further). Mutation enables fine-tuning of the optimal body size, whereas immigration facilitates fitness peak shifts.

### The movement phase

#### Emigration rate

Whether an individual moves depends on the ratio of the amount of energy present within a cell *(R_x,y_)* relative to the maximum amount of energy that can be consumed by all consumers present within that cell. This latter factor is determined by calculating the sum of all individuals’ daily ingestion rates within that cell 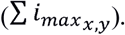

By assuming a symmetric competition, the probability of moving *(p*) is equal for all individuals present within the same cell and is calculated by (based on [44]) :

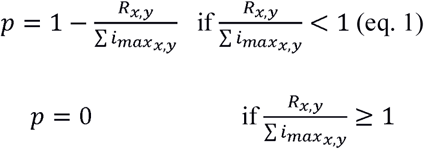

#### Determining cell of destination

As one time step in our model corresponds to one day, we do not model the movement behavior of an individual explicitly but instead, estimate the total area an individual can cover during a day in search for resources. This total area an individual can search during a day is called its foraging area which is circular and is defined by a radius *(rad,* see further). The center of an individual’s foraging area corresponds to its current location. Overall, the size of an individual’s foraging area increases with its size [4,21] and is recalculated daily by taking into account an individual’s optimal speed (Vopt), movement time (t_m_) and perceptual range *(d_per_).* The cost of movement includes the energy invested by an individual in prospecting its foraging area, and is therefore independent of the final cell of destination.

An individual’s average speed of movement (Vopt, in meters per second) is calculated by means of the following allometric equation, derived for walking insects [4,45]:

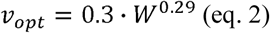

With *W* referring to the weight of an individual in kg, not including the energy stored in its energy reserve. The time an individual invests in movement per day (t_m_ in seconds) is maximally 1 hour. In case too little internally stored energy *(E_r_)* is present to support movement for one hour, *t_m_* is calculated by:

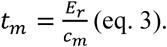

*c_m_* refers to the energetic cost of movement (in joules per second) and is calculated by the following formula, which is based on running poikilotherms [4,45]:

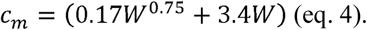

Independent of the cell of destination, the cost of moving during the time *t_m_ (t_m_ c_m_)* is subtracted from an individual’s energy reserve. Based on *tm* and *v_opt_*, the total distance an individual covers at day *t (d_max_)* is determined:

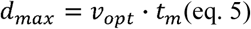

Next, the perceptual range of an individual is determined by means of the following relationship:

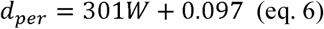

For simplicity, this relationship is linear and based on the assumption that the smallest individual (0.01g) has a perceptual range of 0.10 m and the largest individual (3g) a perceptual range of 1m. The effect of this relationship has been tested (see supplementary material part 4). Moreover, the positive relationship between body size and perceptual range or reaction distance has been illustrated over a wide range of taxa, including arthropods (supplementary information of [46 ]).

The foraging area of an individual is circular and its radius *(rad,* in m) is calculated by taking into account the total distance the individual has covered during the day and the individual’s perceptual range (see supplementary material part 2 for an explanation of this formula):

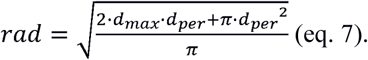

In order to avoid artifacts of applying the continuous variable rad to a grid-based landscape, a random value drawn from the following uniform distribution, [—0.5 · *SL,* 0.5 · *SL],is* added to *rad.*

The selection process for finding a new location within this foraging area depends on the selective ability of the individual. Here, we make a distinction between three types of selection procedures during movement.

#### Case 1- Uninformed movement

Within this scenario, movement is completely uninformed. As such, no distinction can be made between matrix and habitat. Within the foraging area, the new location is randomly sampled.

#### Case 2- Partially informed movement

Here, an individual is able to distinguish matrix from habitat and will always prefer the latter above the former. An individual will sample its location randomly from the suitable cells within its foraging area.

#### Case 3- Informed movement

Here, an individual moves to the cell with the highest amount of resources within its foraging area.

### Immigration

The frequency with which immigrants arrive in the landscape is described by *q.* This variable is fixed at one per 100 days. The process of determining an immigrant’s adults size is similar as during initialization (see below). An immigrant is always introduced within a suitable cell and its energy reserve contains just enough energy to survive the first day.

### Metapopulation and metacommunity perspective

By applying an individual-based approach, we were able to include intra-specific size variation and stochasticity within our model. This approach in conjunction with the assumption of asexual reproduction and equivalent ontogenetic and interspecific scaling exponents [47,48], implies that our results can be interpreted both at the metapopulation and metacommunity level.

### Initialization

Per parameter combination, 10 simulations were run. At the start of a simulation, adult individuals were introduced with an average density of two individuals per suitable cell. The adult weight of each individual (*W*_max_) was determined by drawing the value for log(*W_max_)* from an even distribution between -5 and -2.522878745. Also, each initialized individual carried enough energy within its energy reserve to survive the first day. Initial resource availability per cell corresponded to the maximum carrying capacity. Because of computational limitations, total runtime differed between simulations. For an overview, see supplementary material part 3.

### Data analysis

During each simulation, we traced changes in the mean amount of resources per cell, total number of adults and juveniles, average adult weight *(W_max_)* and the coefficient of variation, skewness, and kurtosis of the consumer’s adult weight *(W_max_)* distribution. Every 500 time steps, the value of *W_max_* of maximum 50 000 randomly sampled individuals was collected.

### Variability

In order to infer the temporal stability of the community at different scales we calculated the *α, β_2_* and *γ* variability for each simulation run. This calculation is based on samples of total consumer biomass every 10 time steps during the final 100 time steps of a simulation within 100 pre-selected, suitable cells. *α* variability is a measure of the local temporal variability and is calculated by

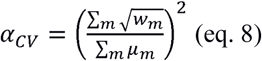

With *w_m_* referring to the temporal variance and *μ_m_* to the temporal mean of population or community consumer biomass in cell *m* [49]. The temporal variability at the metapopulation or metacommunity scale or *γ* variability was calculated by:

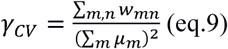

With *w_mn_* referring to the temporal covariance of population or community biomass between cells *m* and *n* [49]. Finally, β_2_?variability or asynchrony-related spatial variability was determined by:

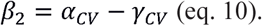

### Reproductive success and movement

Throughout the final 600 days of a simulation, 1000 eggs were randomly selected to be followed during their lifetime. The movements and reproductive success of the resulting consumer individuals were recorded.

### Sensitivity Analysis

A thorough sensitivity analysis was conducted. See supplementary material part 4 for an overview of the tested parameters and their effects.

## Results

A clear interaction with information use is observed when studying the effect of habitat fragmentation and loss on the average body mass of a consumer population or community (Fig 2). Individuals are larger with increasing loss of habitat when movement is fully informed (Fig 2). This effect is enforced by increasing habitat fragmentation (Fig 2). When *P* equals 0.05, *H=0* and movement is informed, 15% of the population does not belong to the smallest size class. Although these larger individuals are lower in abundance than the smallest individuals, they represent a large fraction of total consumer biomass (60%). In contrast, average body mass decreases with habitat fragmentation when movement is uninformed (Fig 2). No clear pattern is observed when movement is partially informed. Still, individuals tend to be smallest within the landscape type with *P* equaling 0.05 and H=1 and small individuals do not occur when *P* equals 0.05 and H=0 (Fig 2 & 3). When comparing body sizes between movement types, individuals with informed movement are the smallest (Fig 2).

**Figure 2:**
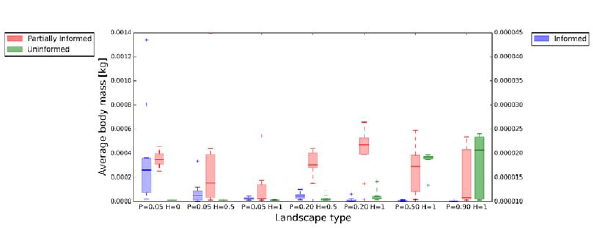
The effect of habitat fragmentation and destruction on average adult body mass of the consumer(s) for each of the three types of information use during movement (informed, partially informed or uninformed). The configuration of suitable habitat within a landscape is described by means of *P* (percentage of suitable habitat) and *H* (level of autocorrelation). Note the different axis scales for partially informed and uninformed movement on the one hand, and informed movement on the other.

The narrowest body size distributions, reflected by the high level of kurtosis, occur in the landscapes with high percentages of suitable habitat (P equaling 0.50 or 0.90) when movement is informed (Fig 3 and S5.4). Overall, most distributions are right-skewed, except for the distributions with partially informed movement, which tend to be neutrally skewed (Fig 3 and S5.11). Because the uninformed and partially informed strategy become identical when *P* approaches one, body size distributions are similar when movement is partially informed or uninformed when *P* equals 0.9 (Fig 3).

**Figure 3:**
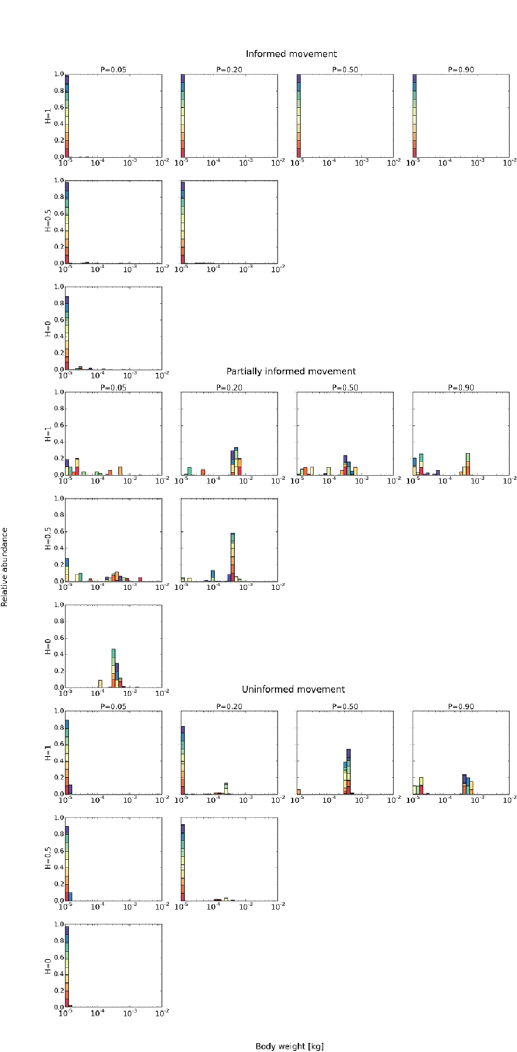
The effect of habitat fragmentation and destruction on the adult body size distribution of a consumer population or community, with movement of the consumer informed, partially informed or uninformed. The configuration of suitable habitat within a landscape is described by *P* (percentage of suitable habitat) and *H* (level of autocorrelation). Each color represents the outcome of a single simulation. In total, ten simulations were run per scenario.

As informed movement results in the selection of the smallest individuals, the highest abundances are observed in these simulations (Fig S5.9). Also, the chance of moving during a day is largest when movement is informed (Fig S5.5). Large individuals can occur in all landscape types when movement is partially informed and in landscapes with a high percentage of suitable habitat when movement is uninformed. Total lifetime is longest in those simulations having the largest individuals (Fig S5.8). As large individuals move further than small individuals (Fig S5.6), their total distance covered during one lifetime is also larger (Fig S5.7).

At the local and inter-patch scale, temporal variability of total consumer biomass is highest when movement is informed (Fig S5.1 and S5.2). However, at the landscape scale, no clear distinction between movement types in temporal variability is observed (Fig S5.3). Still, the landscape type with *P* = 0.05 and *H* = 1 is most variable at the landscape scale when movement is uninformed or partially informed (Fig S5.3). This explains why two out of the ten simulations with partially informed movement went extinct for this landscape type.

Finally, when movement is informed, resource and consumer dynamics at the landscape scale are very stable (Fig S7.5). During a simulation, resources are always spread according to a consistent, homogeneous pattern within the landscape (Fig S7.2). On the contrary, when movement is uninformed or partially informed, resource and consumer dynamics fluctuate strongly in time (Fig S7.1). In addition to these temporal fluctuations, resources are either homogeneously (Fig S7.4) or heterogeneously (Fig 7.3) distributed in space. In some simulations, fluctuations in spatial resource dynamics (homogeneous or heterogeneous spread of resources) correspond with shifts in average size of the consumer (Fig S7.8). However, this is not always the case (Fig S7.9). When movement is informed, resources are only heterogeneously distributed when the landscape is strongly fragmented and contains a low percentage of suitable habitat (Fig S7.6). In this case, resource and consumer dynamics are more unstable (Fig S7.7).

## Discussion

Several theoretical models have investigated how consumer-resource dynamics are affected by nonrandom settlement [39,40], body size distributions [8,14,31,50], spatial habitat configuration [51–54], and more specifically, landscape fragmentation [55]. However, only a few studies combined these research interests in an integrated manner [12,29,30]. Our study is unique as it investigates how body size distributions of a consumer population or community evolve in response to landscape fragmentation and habitat loss, while taking into account the level of informed settlement.

Our model provides a mechanistic understanding of optimal body size distributions and shows that individuals should become larger with increasing fragmentation and loss of habitat when movement is informed, smaller when movement is uninformed and be almost invariant when movement is partially informed. Information use during settlement has a critical impact as it is related to multiple costs during dispersal [56]. When movement is informed, individuals should be able to trace resource availability within the landscape, preventing local resource accumulation. This is in line with our observation that overall, average resource amounts are lowest when movement is informed (Fig S5.10). As such, informed movement results in stable resource amounts and consumer numbers at the landscape scale (Fig S7.5). If resources are homogeneously distributed in space, even small individuals are guaranteed to find resources within their proximity if they are capable of informed movement. As these small individuals have the shortest developmental time, they have a large selective advantage over large individuals and dominate the population when *P* is high. When *P* is low, and especially when *H* is low as well, a small but stable number of large individuals are able to coexist within the landscape as only large individuals are able to trace isolated patches with resources. These patches are out of reach for the smallest individuals, which remain within non-isolated patches. The sensitivity analysis highlights that when the relative mobility of the smallest individuals is decreased, only larger individuals survive when *P* and *H* are low; these findings highlight the role of the trade-off associated with body size with regard to movement (efficiency) and metabolic efficiency. Our finding contradicts that of another theoretical study by Buchmann (2013) [29], who concluded that habitat destruction and fragmentation resulted in a relatively higher frequency of small individuals of mammals and birds. Assuming that mammals’ and birds’ movement is informed, we predict the opposite pattern. This inconsistency may result from differences in model design as their model did not include any resource-consumer dynamics and therefore local colonization-extinction events, which are crucial in shaping body size distributions. Moreover, it did not link body size with developmental time, which drives the selection of small individuals.

On the contrary, when movement is implemented as uninformed or partially informed, individuals do not observe local resource quantity, allowing for resources to accumulate. This results in heterogeneous spatial distributions of the resource. Moreover, resource and consumer dynamics fluctuate strongly in time when movement is not informed (Fig S7.1). When few resources are available within the landscape with large P, there is selection in favor of those individuals that can reach these few patches with resources first (Fig S7.11). Therefore, large individuals can invade the population or community resulting in large-sized equilibria. However, when resources are highly abundant within the landscape, small-sized individuals can reinvade as they have the shortest life cycle and increase fastest in number (Fig S7.10), shifting the equilibrium towards small-sized individuals again. Hence, when *P* is high, a dynamic equilibrium involving two alternative states is observed: one state with small individuals and one state with large individuals [57]. These shifts do not occur when immigration from outside the landscape is turned off, which highlights the significance of immigration as a mechanism maintaining fundamental genetic variance [3]. Some rate of immigration is realistic as open communities are the rule rather than exception in nature [1,58].

When movement is uninformed, individuals decrease in size with decreasing levels of suitable habitat. As large individuals move further, they have the highest chance of ending up outside suitable habitat. This risk is even more elevated when the landscape is less autocorrelated, resulting in even smaller individuals. When *P* equals 0.50 and movement is uninformed, the equilibrium with only small individuals is almost never achieved. Probably, at this particular ratio of suitable versus unsuitable habitat, gaps of unsuitable habitat are relatively easily crossed by large individuals whereas small individuals rarely manage to cross such gaps (see supplementary material part 6). This mechanism might be comparable to the mechanism allowing for emigration-mediated coexistence in food webs: the competitive strength of a strong competitor is lowered by its emigration, enabling coexistence with a weaker competitor [39].

In case of partially informed movement, no clear effect of habitat loss and fragmentation on body size is visible. Still, average body weight is smallest when very few suitable cells are present and they are strongly aggregated (P = 0.05, *H* =1). Consequently, all cells are within reach of the smallest individuals, lowering the advantage of large individuals. Only within this scenario, two out of ten simulations went extinct, indicating that small individuals are vulnerable to extinction under these circumstances. In this scenario, small individuals might go extinct as (i) they have low probability of locating cells with high resource abundance (versus a scenario with informed movement) and (ii) experience strong competition (versus a scenario with uninformed movement). These reasons also explain why small individuals do not occur in any simulation in which the little available habitat is spread widely across the landscape (P = 0.05 and *H* = 0), as then even fewer cells are reachable for the smallest individuals. Therefore, large individuals invade the landscape as they can also access the more isolated cells.

Type of movement not only has a large influence on resource distribution, but also on the spatial structuring of populations or communities. When movement is informed, consumers move more often than when movement is partially informed or uninformed. Movement events can either occur at faster or slower rates than local food web dynamics, resulting in spatially coupled populations (e.g. foraging behavior) or classic metapopulation dynamics (e.g. extinction, colonization events), respectively [59]. As such, we might conclude that patches have a higher tendency of being spatially coupled when movement is informed, than when movement is partially informed or uninformed.

Our sensitivity analyses showed that our model results were robust. Only immigration rate and growth speed of the resource affect the outcome. When the growth speed of the resource and thus productivity is lowered, no large individuals are observed in any simulation and many simulations go extinct. As large individuals need a minimum amount of resources to survive, they are no longer able to persist. When immigration rate is deactivated, large individuals completely disappear in some scenarios (e.g.when *P =* 0.90, *H* = 1 and movement is uninformed) as they occur at much lower abundances than small individuals and are therefore more susceptible to drift. However, when large individuals remain in a certain scenario without immigration, the strength of selection in favor of these large individuals is illustrated.

## Conclusions

Empirical inconsistencies in body size responses to habitat loss and fragmentation have so far been attributed to differences in scale [22] and in the suitability of the matrix [27] and whether an equilibrium was obtained (e.g. extinction time lags) [20]. Our model provides an alternative explanation: the level of informed movement. Moreover, it highlights the relevance of not only habitat loss but also of fragmentation, since the latter reinforces the effect of the former. Importantly, our model reveals that habitat fragmentation and loss lead to a possible introduction of large individuals or species when settlement is informed and a disappearance of large individuals when settlement is uninformed. Our results are of great relevance to conservation management. Not only body size distributions are affected by habitat fragmentation but also the distribution of resources (changing from homogeneous to heterogeneous) and stability of consumer-resource dynamics (from stable to unstable), implying an elevated extinction risk.

## Authors’ contributions

DB, TH and JH conceived the ideas and designed methodology; JH designed the model; DB, MLV, TH and JH analyzed the data; DB, MLV and JH led the writing of the manuscript.

